# Decoding the Glycan Signature: Unraveling *N*-Glycosylation Alterations in Glycogen Storage Disease Ia and Ib

**DOI:** 10.1101/2025.10.23.684118

**Authors:** Ruiqi Xiao, Jing Zheng, Hector F. B. R. Loponte, Yajie Ding, Candelas Gross-Valle, Terry G.J. Derks, Peter L. Horvatovich, M. Rebecca Heiner-Fokkema, Barbara M. Bakker, Justina C. Wolters, Guinevere S. M. Lageveen-Kammeijer

## Abstract

Glycogen storage disease (GSD) types Ia and Ib are rare inherited metabolic disorders caused by pathogenic variants in *G6PC* or *SLC37A4*, respectively. These defects disrupt glucose homeostasis and may affect protein glycosylation. We systematically profiled sera *N*-glycomes from retrospectively collected GSD Ia (n=17), GSD Ib (n=8), and control (n=21) samples. Derived traits analyses revealed distinct, subtype-specific, and shared glycomic alterations, including shifts from α2,3-to α2,6-sialylation in both subtypes. GSD Ia showed enhanced branching and reduced fucosylation, whereas GSD Ib displayed elevated fucosylation, bisection, and reduced branching. In GSD Ia patients with hepatocellular adenoma/carcinoma (HCA/HCC), further enrichment of oligomannosidic and α2,3-sialylation was observed. Several individual *N*-glycans showed strong discriminatory performance, supporting their potential as biomarkers for GSD I subtyping and HCA/HCC surveillance. This study provides the first comprehensive characterization of systemic *N*-glycans in GSD Ia and Ib, revealing glycomic remodeling as a potential biomarker of disease subtypes and tumor progression.

**Take-home message (synopsis) of the article:** Serum/plasma *N*-glycomic profiling reveals distinct glycosylation signatures in GSD Ia and GSD Ib, including tumor-associated shifts in GSD Ia, highlighting novel biomarkers for disease subtyping and surveillance of hepatic complications.

## 1. Main

Glycogen storage diseases (GSDs) are rare, severe inherited metabolic diseases caused by impaired glycogen metabolism.^1^ Among them, GSD type I (GSD I) is one of the most severe, with a prevalence of approximately 1 in 100,000 live births.^2^ GSD I comprises subtypes Ia (OMIM:#232200; ORPHA:79258) and Ib (OMIM:#232220; ORPHA:79259), which result from pathogenic variants in the *G6PC1* ^3^ and *SLC37A4* ^4^ genes, respectively (**Figure 1A**). These genes encode glucose-6-phosphatase (G6PC1; type Ia) and the glucose-6-phosphate transporter (G6PT; type Ib), both localized to the endoplasmic reticulum (ER) membrane. ^5, 6^ Under physiological conditions, G6PT imports glucose 6-phosphate (G6P) from the cytoplasm into the ER, where G6PC1 hydrolyzes it into free glucose and inorganic phosphate. Subsequently, the glucose is transported into the bloodstream and thereby contributes to systemic glucose homeostasis (**Figure 1A**).^7–9^

**Figure 1.**
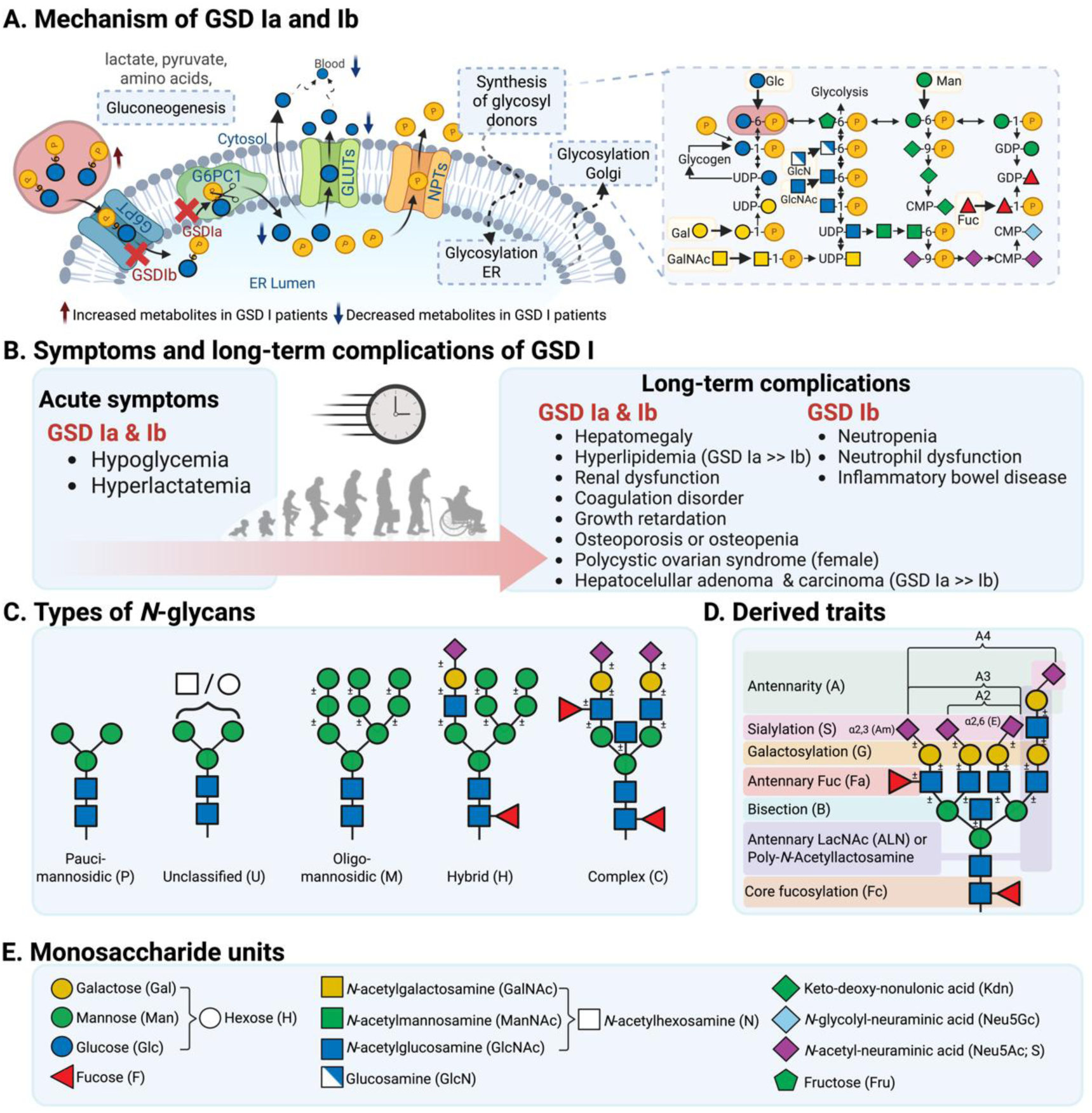
Overview of GSD I mechanisms, clinical manifestations, and glycomic analysis. **(A)** Schematic representation of the defective glucose-6-phosphatase (G6Pase) system in GSD Ia and the G6P transporter (G6PT) in GSD Ib, highlighting their respective roles in hepatic glucose metabolism and the resulting metabolic block. The inset illustrates the biosynthesis of nucleotide sugar donors in the cytoplasm, which are essential substrates for *N*-glycan assembly in the endoplasmic reticulum (ER) and Golgi apparatus. **(B)** Immediate clinical symptoms and long-term complications associated with GSD I, including hypoglycemia, hepatomegaly, etc., and a higher risk of hepatocellular adenomas (HCA) and carcinoma (HCC) in GSD Ia than in GSD Ib. **(C)** Representative structures of major *N*-glycan types— paucimannose, oligomannose, hybrid, and complex— are shown, classified according to branching and monosaccharide composition. *N*-glycans placed in the unclassified group are those that do not fully meet the criteria defining any of the established classes. **(D)** Derived glycan traits grouped by biosynthetic features such as antennarity, fucosylation, sialylation, and galactosylation, enabling comparison of pathway-related alterations in glycosylation. **(E)** Monosaccharide symbols and linkages used throughout the figures, following standard glycan notation. A yellow circle with a P inserted indicates phosphate.

In GSD Ia, deficiency of G6PC1 causes accumulation of G6P in the ER and reduced glucose release into the circulation,^3,8^ whereas in GSD Ib, defective G6PT prevents G6P transport into the ER, leading to depletion of both G6P and glucose within the ER lumen, as well as reduced glucose release into the circulation.^4,8^ The *G6PC1* gene is mainly expressed in the liver, kidney, and intestine,^1,3,5^ whereas *SLC37A4* shows a broad expression across multiple tissues, including liver, kidney, pancreas, intestine, and immune cells.^4,6^ The metabolic disturbances result in hypoglycemia, hyperlactatemia, and hyperlipidemia,^8^ along with long-term complications, including liver damage, kidney failure, and hormone disorders.^3, 10, 11^ GSD Ia patients are particularly prone to developing hepatocellular adenoma (HCA) and hepatocellular carcinoma (HCC),^12^ whereas GSD Ib patients characteristically develop neutropenia (**Figure 1B**).^13^ Despite these distinct phenotypes, there is lack of reliable biomarkers to assess the risk of developing these complications. In addition, our recent proteomics study suggested small protein differences and pathway changes between GSD Ia and Ib, including several proteins involved in glycoprotein changes. ^14^ Moreover, the glycan synthesis is tightly linked to glucose metabolism. Therefore, investigating protein *N*-glycosylation could provide valuable novel biomarkers. G6P not only serves as a key intermediary between glycolysis and glycogen metabolism but also provides the precursor for nucleotide sugar donors required for *N*-glycan biosynthesis (**Figure 1A**). Consequently, differences in G6P availability in cellular compartments between GSD Ia and Ib may alter the supply of glycosylation precursors, thereby driving subtype-specific *N*-glycan profiles.

*N*-glycosylation is a critical post-translational modification that ensures proper protein folding, stability, and function, ^15^ impacting cellular signaling ^16, 17^ and immune recognition^18,19^. The *N*-glycan synthesis begins in the ER with the assembly of a lipid-linked oligosaccharide precursor, its transfer to the target protein, followed by subsequent stepwise trimming and modifications in the ER and Golgi apparatus.^20^ This process is tightly regulated by the availability of glycosyl donors, which are derived from intracellular and intracompartmental G6P levels.^15,20,21^ Structurally, *N-* glycans are classified into pauci-mannosidic (P), oligo-mannosidic (M), hybrid (Hy), complex (C), and unclassified (U) types (**Figure 1C**). ^22^ To extract biologically meaningful features across this structural diversity, derived glycan traits can be assigned, reflecting properties such as antennarity (A), sialylation (S), and fucosylation (F) (**Figure 1D**).^22^ The monosaccharide units (**Figure 1E**) form the structural building blocks underlying these modifications.

Previous studies on multiple GSD subtypes suggest that GSDs are associated with altered protein glycosylation, in some cases resembling congenital disorders of glycosylation (CDG). For example, in GSD XIV, truncated transferrin *N*-glycans were linked to reduced uridine diphosphate-galactose (UDP-Gal) levels. ^23^ Truncated complex-type *N*-glycans have also been described in neutrophils of GSD Ib and G6PC3-deficient patients. ^24^ In serum from GSD Ib patients, elevated high mannose and hybrid-type *N*-glycans were reported,^25^ while in GSD types III and IX, aberrant ApoC-III glycosylation was identified.^26^ Together, these findings underscore the importance of protein glycosylation defects in GSD, yet comprehensive *N*-glycomic profiling of GSD Ia and Ib has not been reported.

In this retrospective, non-interventional cohort study, we systematically profiled the serum and plasma *N*-glycomes of GSD Ia and Ib patients using high-sensitivity capillary electrophoresis-electrospray ionization mass spectrometry (CE-ESI-MS). ^27^ We explored whether disruptions in G6P metabolism give rise to distinct glycosylation signatures and whether these alterations can serve as biomarkers for disease subtyping and tumor surveillance. By linking metabolic defects to glycomic changes, this work addresses a critical gap in understanding the molecular basis of GSD I complications and explores the potential of *N*-glycans as prognostic biomarkers.

## 2. Results

In this study, we investigated whether disruptions in glucose-6-phosphate metabolism in GSD Ia and Ib patients affect circulating protein *N*-glycosylation. Using high-sensitivity CE-ESI-MS, we systematically profiled serum and plasma *N*-glycans to reveal disease-associated alterations and to identify potential biomarkers for monitoring disease progression and complications. We first examined global *N*-glycome differences, then explored shared, opposing, and subtype-specific derived traits, and finally assessed individual glycans for their biomarker potential, including in patients with hepatic tumors.

### Global N-glycome differences between GSD Ia, GSD Ib, and controls

We identified 249 unique *N*-glycan compositions (detected in >50% of samples in at least one group; **Supplementary Information, Table S1; Figure S1A**). All the masses of their precursor ions were confirmed based on accurate mass (± 10 ppm), isotopic pattern (< 20% deviation), and migration behavior. Of these, 215 glycan structurally confirmed by MS2, with 80 *N*-glycan compositions were supported with fragmentation spectra, and 135 glycan compositions confirmation was based on accurate mass (< ± 10 ppm), isotopic pattern (< 20% deviation), and migration behaviour. In addition, 34 *N*-glycans showed ambiguities in their compositions and could not be fully resolved; these were included in the individual *N*-glycan analysis, but were excluded from the composition-specific derived trait analysis (**Figure 1C** and **1D; Supplementary Information, Table S1**). Minimal technical variation was demonstrated by the clustering of total plasma *N*-glycome (TPNG) standard sample replicates (n=5) in the principal component analysis (PCA) (**Supplementary Information, Figure S1; Table S2**). This indicates that the differences observed in the GSD I patient samples reflect true biological variation rather than technical noise.

PCA of the total *N*-glycans revealed clear separation between GSD Ia, GSD Ib, and control samples, with 14.3% and 12.9% of the variance (**Supplementary Information, Figure S2B**). No clustering was observed for potential confounders such as age, sex, sample type, feeding state, or HCA/HCC status (**Supplementary Information, Figure S3**). Volcano plots showed that 82 (38%), 37 (17%), and 56 (26%) *N*-glycan compositions were significantly altered in GSD Ia vs. controls, GSD Ib vs. controls, and GSD Ia vs. Ib, respectively (**Supplementary Information, Figures S2C-S2E; Table S3**). These findings highlight pronounced differences in circulating *N-*glycans across GSD subtypes and controls.

### Derived traits reveal shared, opposing, and subtype-specific alterations

To capture functionally related glycosylation changes, we derived composite traits from 215 structure-confirmed *N-*glycans (**Supplementary Information, Table S4**). Based on biological pathways, these derived traits were grouped according to shared structural characteristics, such as antennarity, sialylation, and fucosylation (**Figure 1D**), thereby providing a more integrated view of the disease-associated alterations. PCA plot of derived traits showed partial separation between GSD Ia, GSD Ib, and controls (**Figure 2**, pairwise comparisons in **Supplementary Information, Figure S4A-C**). The 20 most affected traits were mapped onto the PCA plot, which revealed three categories of changes, namely traits common to both GSD subtypes compared with controls (**Figure 3**), traits with opposing effects between GSD Ia and Ib (**Figure 4**), and subtype-specific alterations relative to controls (**Figure 5**).

**Figure 2.**
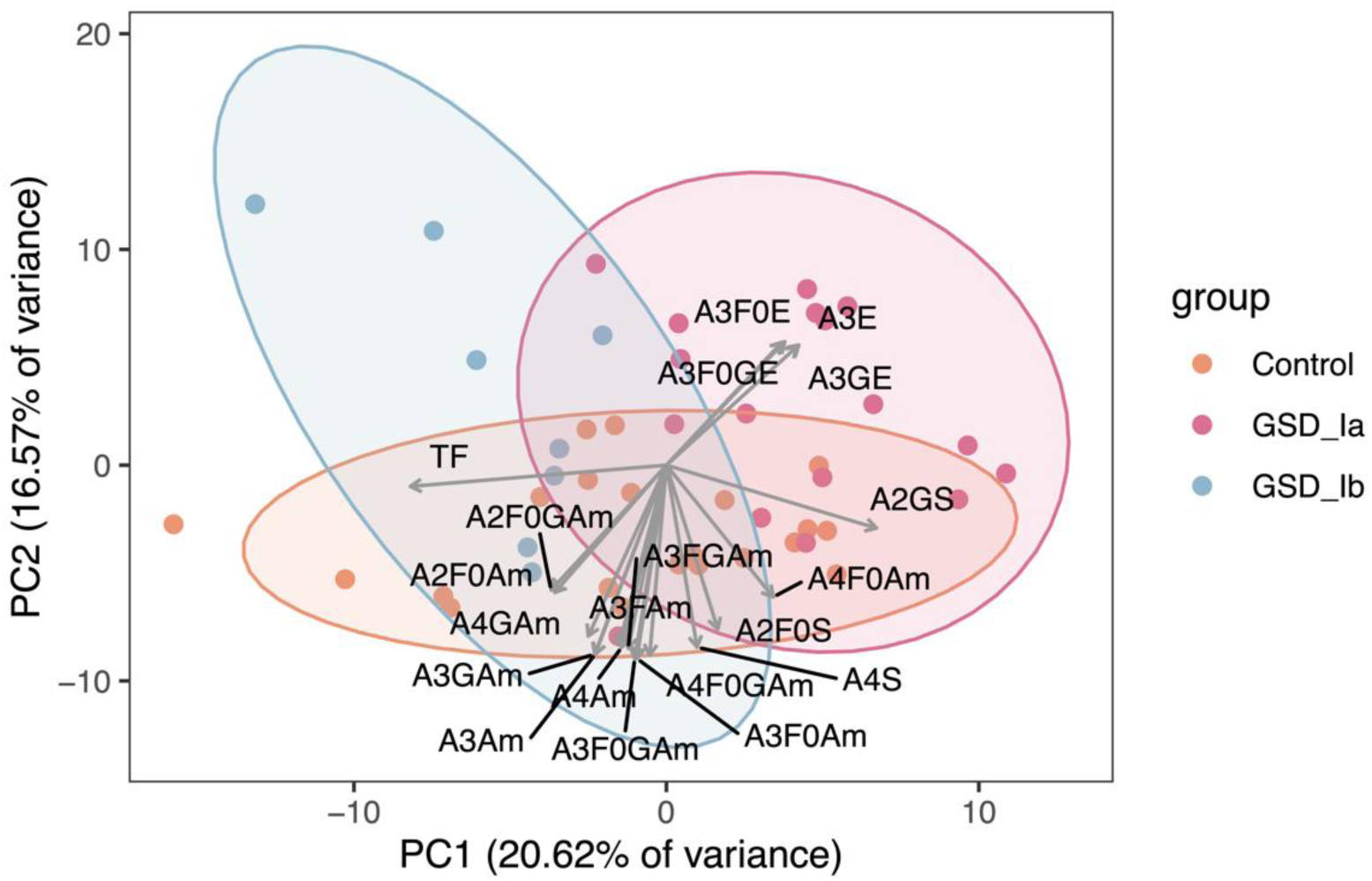
Principal component analysis (PCA) plot of derived *N*-glycan traits. The distribution of samples was based on GSD Ia, GSD Ib, and control groups. Arrows indicate variables that significantly influence the distribution (p < 0.05). The direction of each arrow shows the correlation with the principal components, and the length reflects the strength of the correlation. Example of the derived trait labels: “TF” refers to total fucose. “A3F0E” refers to nonfucosylated α2,6-sialylated triantennary *N*-glycans.

**Figure 3.**
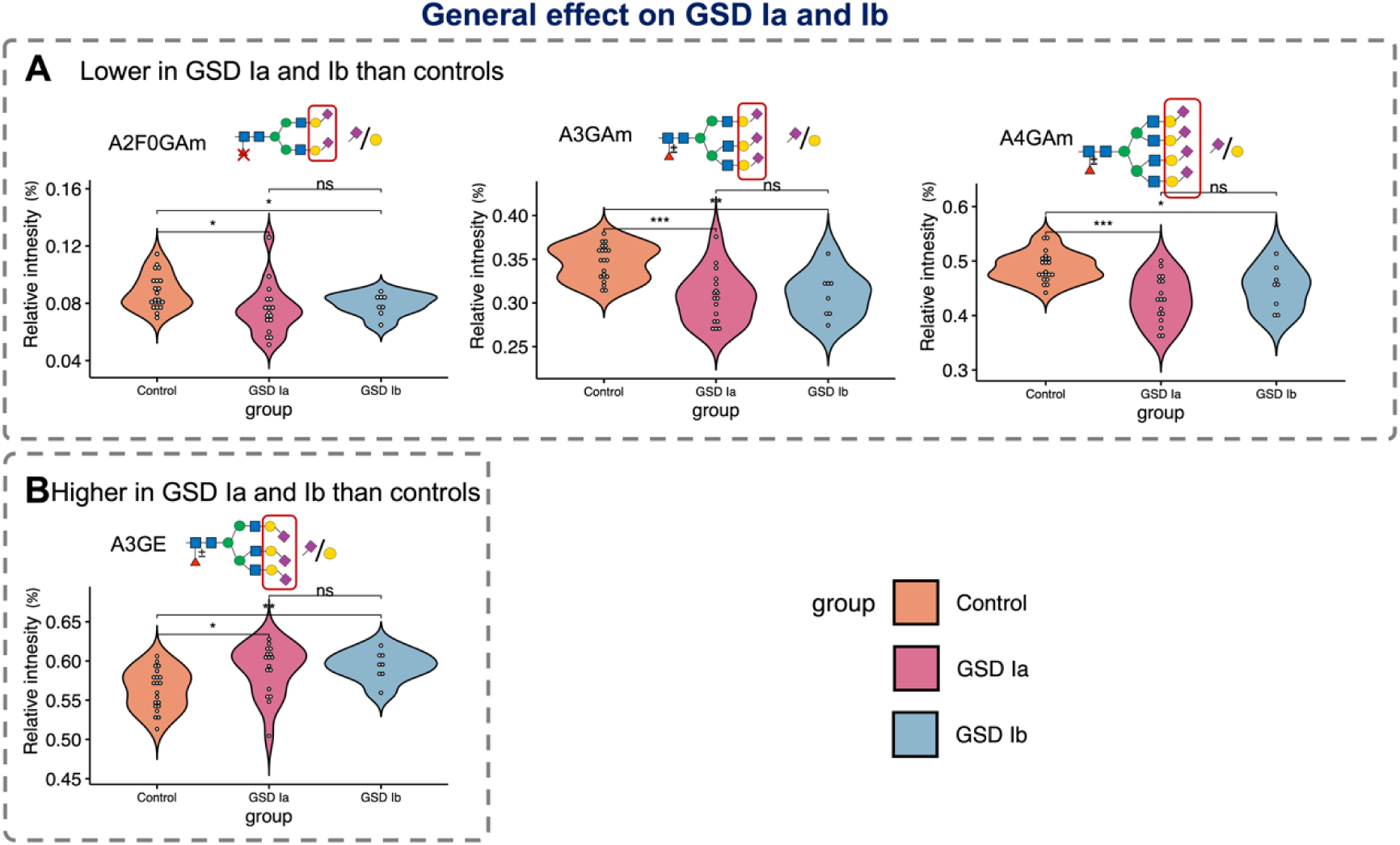
Violin plots of the derived traits with the general effect on GSD Ia and Ib. **(A)** Selected example *N*-glycan traits (**A**) with a lower intensity and (**B**) a higher intensity in GSD Ia and Ib than in control samples (additional *N*-glycan traits can be found in **Supplementary Information, Figure S5**). Significance levels: FDR<0.05 (*), FDR<0.01 (**), FDR<0.001 (***). “ns” refers to no significant changes. Example of the derived trait labels: “A2F0Am” refers to nonfucosylated α2,3-sialylated diantennary *N*-glycans. “A2F0GAm” refers to the ratio of α2,3-sialic acid and galactose on nonfucosylated α2,3-sialylated diantennary *N*-glycans. “A4Am” refers to α2,3-sialylated tetra-antennary *N*-glycans. “A3E” refers to α2,6-sialylated triantennary *N*-glycans.

**Figure 4.**
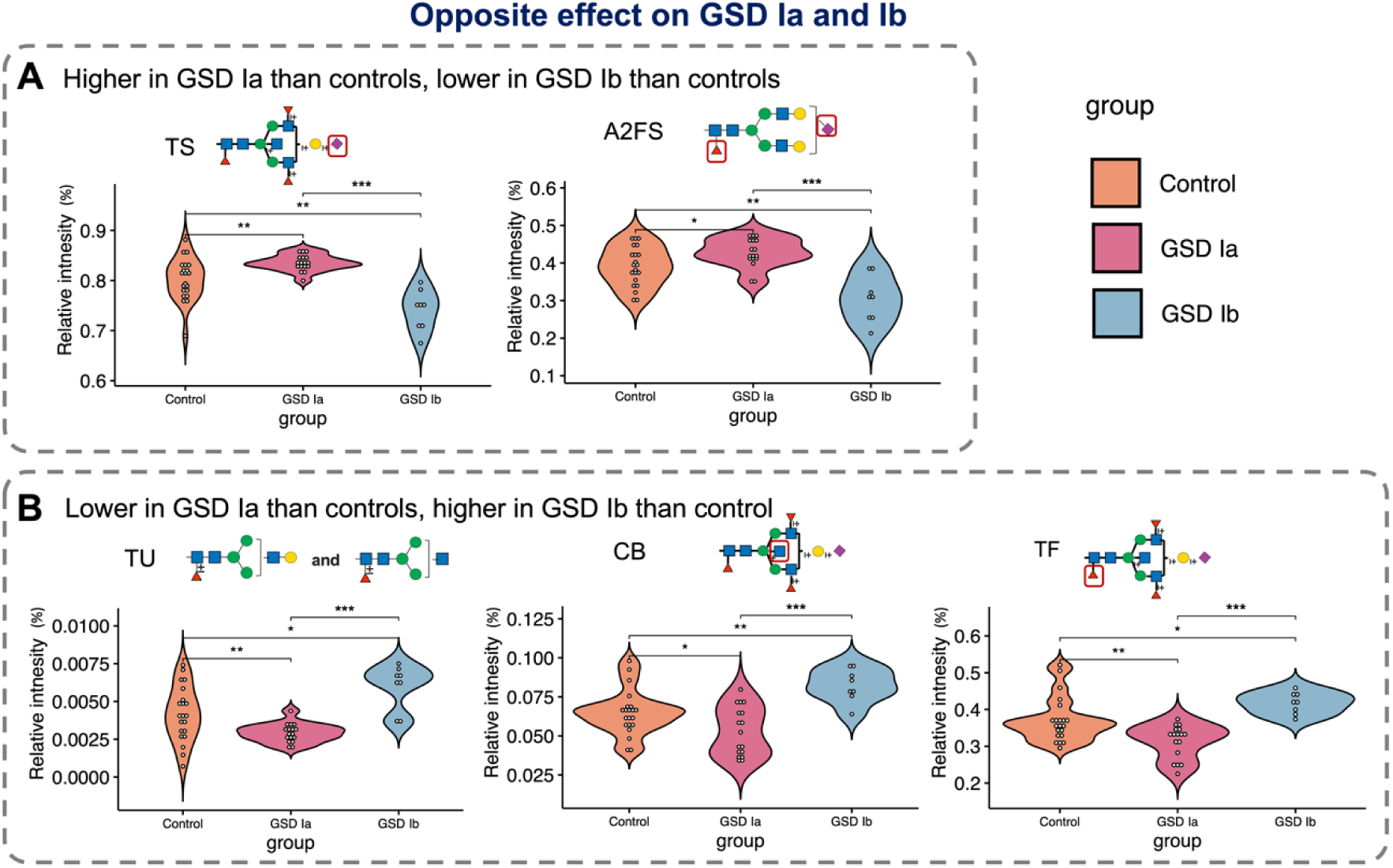
Violin plots of the derived traits with the opposite effect on GSD Ia and Ib compared to control. Compared with controls, **(A)** glycans with a higher intensity in GSD Ia, and lower intensity in GSD Ib, as well as **(B)** glycans with a lower intensity in GSD Ia, and higher intensity in GSD Ib. Significance levels: FDR<0.05 (*), FDR<0.01 (**), FDR<0.001 (***). Example of the derived trait labels: “TU” refers to total unclassified glycans. “CB” refers to bisection complex type *N*-glycans.

**Figure 5.**
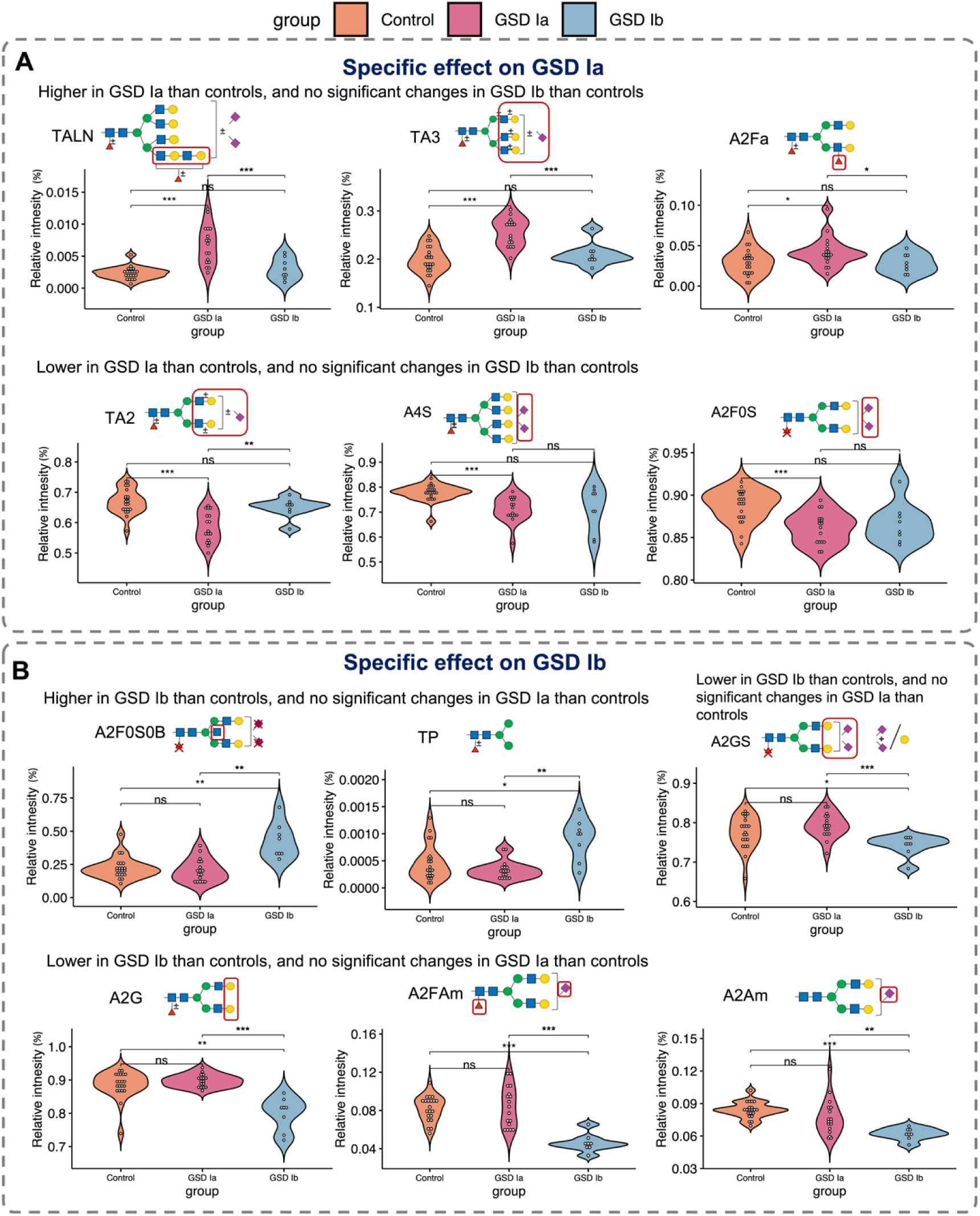
Violin plots of the derived traits with the specific effect on GSD Ia or Ib. **(A)** Glycans with a higher or lower intensity in GSD Ia than controls, but with no significant changes in GSD Ib than controls. **(B)** Glycans with a higher or lower intensity in GSD Ib than controls, but with no significant changes in GSD Ia than controls. Significance levels: FDR<0.05 (*), FDR<0.01 (**), FDR<0.001 (***). “ns” refers to no significant changes. Example of the derived trait labels: “TALN” refers to total *N*-glycans with antennary LacNAc (ALN). “A2Fa” refers to antennary fucosylated diantennary *N*-glycans. “TP” refers to total pauci-mannosidic *N*-glycans.

*Features that are shared between GSD Ia and Ib.* Both GSD Ia and Ib displayed reduced α2,3-sialylation (Am), independent of antennarity or fucosylation, and increased triantennary α2,6-sialylation (E), consistent across PCA and univariate analysis (**Figures 2-3**; **Supplementary Information, Figure S5**).

*Opposing features.* Five traits showed opposite patterns when comparing GSD Ia and Ib (**Figure 4**). In GSD Ia, sialylation level was increased relative to control, whereas in GSD Ib it was decreased, including total sialylation (TS) and sialylation per fucosylated diantennary structures (A2FS) (**Figure 4A; Supplementary Information, Figure S4B**). In contrast, bisection (CB) and total fucosylation (TF) were reduced in GSD Ia but elevated in GSD Ib (**Figure 4B; Supplementary Information, Figure S4B-C**). Similarly, unclassified *N*-glycans (TU), comprising four simple *N*-glycan compositions (H3N3, H3N3F1, H4N3, and H4N3F1) were decreased in GSD Ia but increased in GSD Ib.

*Sub-type specific features.* GSD Ia was characterized by increased antennarity, including triantenary (A3) and tetraantennary (A4) with poly-*N*-acetyllactosamine (ALN, i.e. LacNAc) structures, as well as diantennary *N-*glycans with antennary fucosylation (A2Fa) (**Figure 5A**; **Supplementary Information, Figure S4A**). In contrast, diantennary (A2), sialylation (both α2,3 and α2,6) of tetra-antennary (A4S), and nonfucosylated diantennary (A2F0S) *N*-glycans were specifically decreased in GSD Ia (**Figures 2** and **5A**). In GSD Ib, subtype-specific changes were mainly characterized by increased (bisected) diantennary structures lacking sialylation (A2F0S0B) and total pauci-mannosidic *N*-glycans (TP), and decreased galactosylation (A2G) and sialylation (A2GS, A2FAm, and A2Am) on diantennary *N*-glycans (**Figure 5B; Supplementary Information, Figure 4B**).

### HCA/HCC patients show distinct N-glycan alterations

Eight serum samples in the GSD Ia cohort were from four patients with hepatic tumors (seven HCA, one HCC), which were confirmed by clinical and histological evaluation. The serum *N*-glycans from these patients showed distinct glycosylation changes compared to both healthy controls and GSD Ia patients without tumors. Although PCA did not reveal distinct clustering of patients with or without tumors (**Supplementary Information, Figure S3F**), trait-level analysis identified distinct tumor-associated glycosylation changes (**Supplementary Information, Figures S4D-E** and **Table S4**). These included a pronounced decrease in complex-type structures (TC), enrichment of oligomannosidic (TM) *N*-glycans, both with FDR < 0.001, and selective increases in α2,3-sialylation of fucosylated diantennary *N*-glycans (A2FAm and A2FGAm) (**Figure 6A-B**).

**Figure 6.**
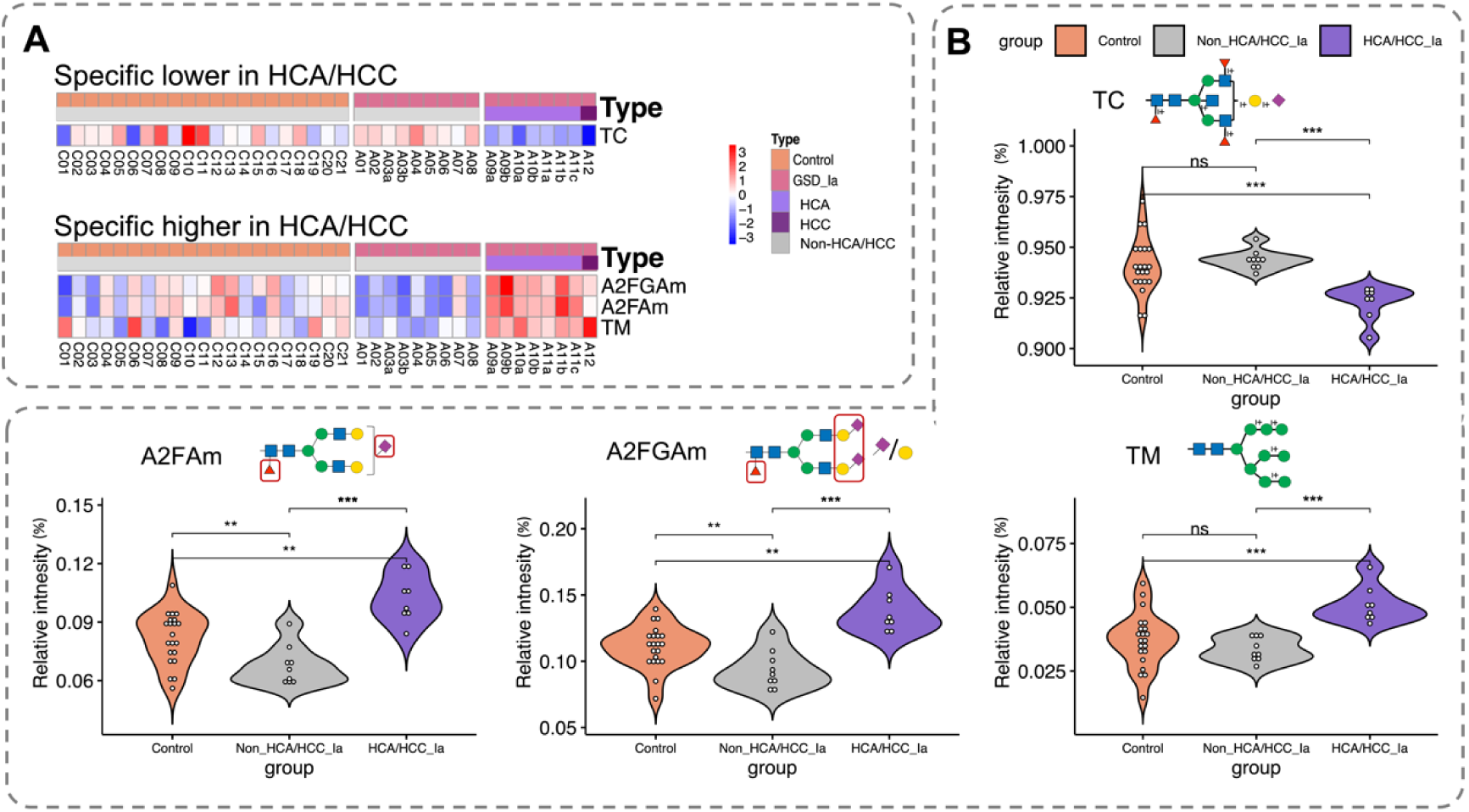
Significantly changed derived traits in GSD Ia with HCA/HCC. (**A**) Heatmap showing the z-score and (**B**) violin plots of these derived *N*-glycan traits specifically in GSD Ia with HCA/HCC. (**B**) Significance levels: FDR<0.05 (*), FDR<0.01 (**), FDR<0.001 (***). “ns” refers to no significant changes. “TC” refers to total complex type *N*-glycans. “A2FAm” refers to fucosylated α2,3-sialylated diantennary *N*-glycans. “A2FGAm” refers to the ratio of r. “TM” refers to total oligo-mannosidic *N*-glycans.

### Individual N-glycans emerge as biomarker candidates

To evaluate translational potential, we next examined individual *N-*glycans, which are more amenable to targeted detection as potential biomarker candidates (**Supplementary Information, Table S3**). Significantly changed *N*-glycans with log_2_^FC^ exceeding ± 2 in pairwise comparisons (**Supplementary Information, Figures S2C-E**) were considered candidates (**Figure 7A**; ROC curves in **Supplementary Information, Figure S6**).

**Figure 7.**
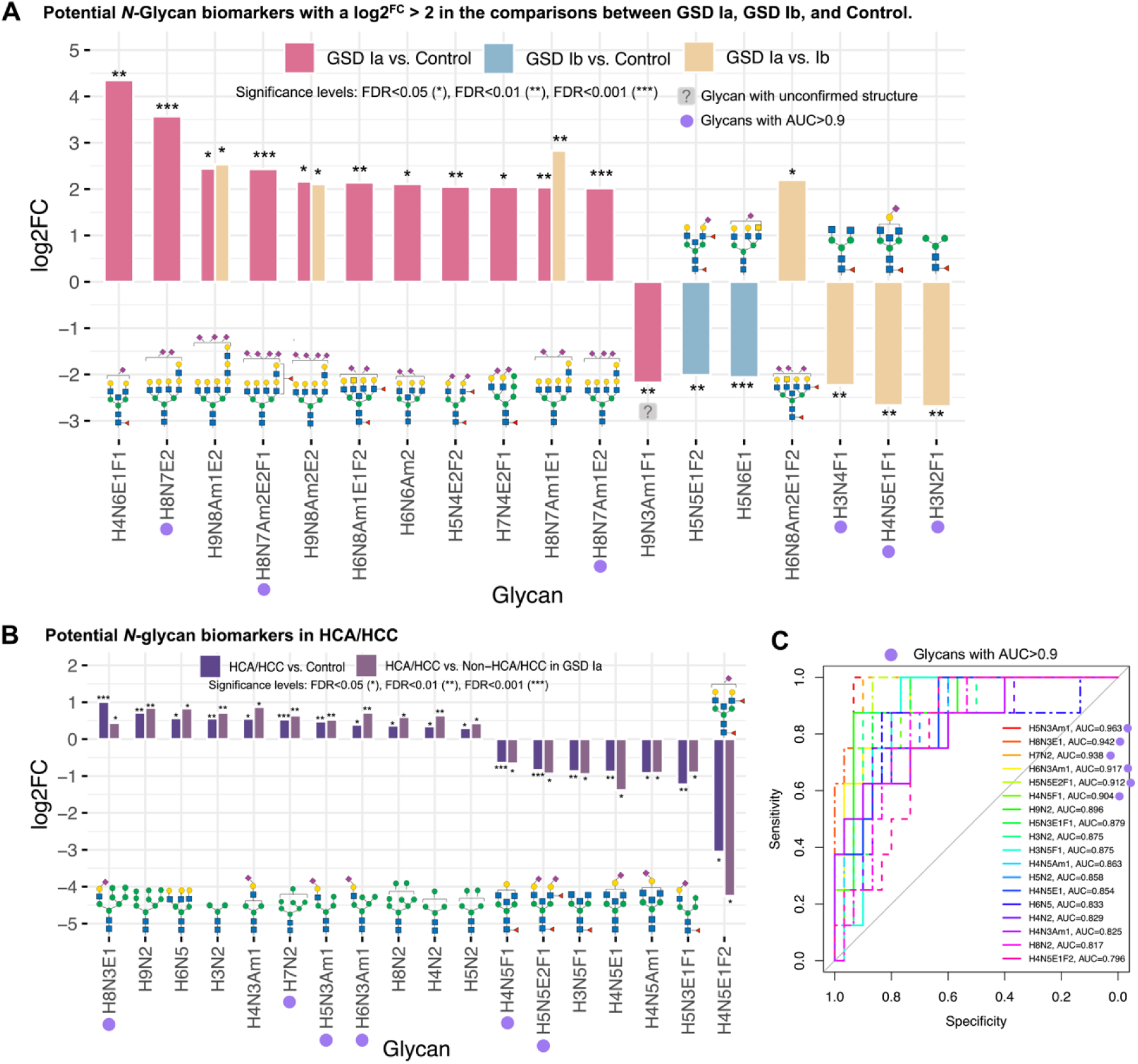
Summary of potential *N-*glycan biomarkers of GSD Ia, GSD Ib, and HCA/HCC. **(A)** Boxplots showing the *N*-glycans with log2^FC^>2 in GSD Ia vs. control, GSD Ib vs. control, and GSD Ia vs. Ib. **(B)** Boxplots showing the Log2^FC^ of *N*-glycans with the same changes in HCA/HCC versus control and Non-HCA/HCC in GSD Ia (n=9). **(C)** Receiver operating characteristic (ROC) curve of *N*-glycans showing in B.

*GSD Ia biomarkers*. A total of 12 *N*-glycans differentiated GSD Ia from controls, including six ALN-rich tetra-antennary *N*-glycans, consistent with the TALN shift toward complex and highly branched structures (**Figure 5A**). Three of these (H8N7Am1E2, H8N7Am2E2F1, H8N7E2) achieved excellent classification performance with AUCs > 0.90, while the AUCs of the remaining three (H9N8Am1E2, H9N8Am2E2, and H8N7Am1E1) were slightly lower (>0.77) (**Figure 7A; Supplementary Information, Figure S6A**), but these later three differentiated GSD Ia not only from controls but also from GSD Ib (**Figure 7A; Supplementary Information, Figure S6C-D**).

*GSD Ib biomarkers.* Two simpler diantennary *N*-glycans (H5N5E1F2 and H5N6E1) were decreased in GSD Ib (log_2_^FC^ < -2), with AUCs > 0.79 (**Figure 7A; Supplementary Information, Figure S6B**).

*HCA/HCC biomarkers.* In GSD Ia with HCA/HCC, a total of 56 and 29 individual *N-* glycans differed relative to controls and non-tumor GSD Ia, respectively, with 18 overlapping between them (**Supplementary Information, Figure S7**). Six pauci-/oligomannosidic *N*-glycans (H3N2, H4N2, H5N2, H7N2, H8N2, and H9N2) were consistently increased, while six bisected *N*-glycans (H4N5F1, H5N5E2F1, H3N5F1, H4N5E1, H4N5Am1, and H4N5E1F2) were decreased (**Figure 7B**). Six *N*-glycans (H5N3Am1, H8N3E1, H7N2, H6N3Am1, H5N5E2F1, and H4N5F1) achieved excellent classification performance (AUCs > 0.90) for separating patients with or without tumors (**Figure 7C**).

## 3. Discussion

This study provides the first systematic *N-*glycomic profiling of GSD Ia and Ib, revealing both shared and subtype-specific glycosylation abnormalities. By linking disruptions in glucose-6-phosphate metabolism to alterations in protein *N*-glycosylation, we demonstrate that glycomic signatures not only distinguish GSD Ia versus Ib but may also serve as candidate biomarkers for disease monitoring and the prognosis of liver adenoma and carcinoma.

The analyses identified consistent alterations across major glycosylation pathways, with sialylation emerging as a unifying feature in both subtypes, and divergent changes in antennarity and fucosylation defining subtype-specific patterns. In GSD Ia with HCA/HCC, we further observed tumor-associated remodeling of the glycome, consistent with glycosylation signatures reported in other cancers. Finally, several individual and composite glycans displayed strong discriminatory performance, underscoring their translational potential as candidate biomarkers. In the following sections, we discuss these findings in relation to sialylation, antennarity and fucosylation, tumor-associated glycosylation, and biomarker interpretation, before outlining the limitations and future directions.

Sialylation emerged as the most prominent alteration in both GSD I subtypes compared with controls. Both GSD Ia and Ib showed a general reduction in α2,3-sialylation alongside an increase in triantennary α2,6-sialylation. To our knowledge, this is the first study to differentiate these linkage-specific changes in GSD I. As terminal residues of the glycan chain, sialic acids contribute to protein stability and act as the molecular recognition sites that protect glycoproteins from premature clearance. ^28-30^ They interact with sialic-acid-binding immunoglobulin-like lectins (Siglecs) and selectins, thereby shaping immune recognition and modulating inflammation.^31–33^ In general, α2,3-linked sialylation is linked to cell adhesion and pro-inflammatory signaling, ^34^ whereas α2,6-linked sialylation has been implicated in immune tolerance and anti-inflammatory regulation. ^35, 36^ In our previous liver proteome analysis of the same GSD I patients,^14^ we observed a decrease in the key enzyme ST6GAL1 (**Supplementary Information, Figure S8; Table S5**), which is an α2,6-sialyltransferase that catalyzes α2,6-sialylation of terminal galactose (**Supplementary Information, Table S6**).^37^ This finding points to a more complex relationship between the hepatic enzyme abundance and the observed glycosylation patterns in circulating glycoproteins. In addition, ST6GAL1 is expressed in many other tissues, including lymphoid organs, pancreatic β-cells, renal tubular cells, and gastric epithelial cells,^38–41^ where it may influence systemic glycosylation patterns. This broader tissue distribution could provide an alternative rationale for the lack of a similar response between the hepatic ST6GAL1 abundance and the circulating glycosylation changes.

When examining subtype-specific changes in sialylation, we detected a decreased total sialylation (TS, A2FS, A2GS, and A2Am) specifically in GSD Ib, consistent with an earlier report of mild hyposialylation of ApoC-III in both subtypes.^26^ In contrast, we found an increase in total sialylated *N*-glycans (TS) in GSD Ia with selective decreases in sialylated tetra-antennary *N*-glycans (A4S) and nonfucosylated sialylated diantennary *N*-glycans (A2F0S). These alterations may reflect the broader scope of our analyses, which covered the total serum/plasma *N*-glycome, whereas the previous study focused specifically on APOC-III, a relatively low-abundant plasma protein (∼0.1-0.2% of the total plasma proteome^42,43^).

Clinically, the observed shifts in sialylation provide insight into the different pathophysiologies of the two GSD I subtypes. Hypersialylation of tumor cell surfaces is a recognized hallmark of many malignancies,^44–46^ promoting immune evasion through Siglec binding, ^30,31,33^ suppression of natural killer (NK) cells and T lymphocytes activity, and macrophage polarization.^44^ The hypersialylation observed in GSD Ia may therefore contribute to a pro-tumorigenic environment and contribute to explain the elevated risk of HCA/HCC in this subtype. In contrast, hyposialylation may impair immune surveillance by reducing molecular recognition and enhancing abnormal protein clearance,^47,48^ which potentially contributes to the recurrent inflammatory phenotype in GSD Ib patients. Together, these findings improve our understanding in the protein glycosylation changes and underscore protein sialylation as a key mechanism affected in GSD I, with distinct differences between GSD Ia and Ib.

Beyond sialylation, we identified distinct subtype-specific changes in antennarity and fucosylation. Compared to controls, GSD Ia exhibited increased tri- and tetra-antennary structures and enhanced ALN, whereas GSD Ib shifted toward simpler pauci-/ oligomannosidic *N*-glycans (TP, TU). Our previous liver proteome analysis partly supports these findings with the specific elevation in GSD Ib for MGAT2 (**Supplementary Information, Figure S8; Table S5**),^14^ which extends biantennary branching (**Supplementary Information, Table S6**).^49^ It will be interesting to further study how this is affected by the observed decrease of the enzymes required for trimming and initiating branching (MGAT1, MAN1A1, and MAN1A2, **Supplementary Information, Figure S8; Tables S5 and S6**),^14,50-52^ which were decreased in both subtypes. It would furthermore be useful to also get information about the enzymes MGAT4 and 5, which are involved in the linkage of the tri/tetra antennary structure (**Supplementary Information, Table S6**), which were not detected in the liver proteome study.

In GSD Ib, galactosylated diantennary glycans (A2G) were decreased compared with controls. This is in line with earlier reports of neutrophil hypo-galactosylation.^24^ We can now link this hypo-glycosylation specific to the diantennary *N*-glycans. Bisection pattern changes revealed further subtype specificity: GSD Ia showed reduced bisection (CB), whereas GSD Ib exhibited increased bisection (CB and A2F0S0B) compared with controls. Because bisection sterically hinders further branching, its decrease in GSD Ia may promote the observed rise in ALN and branching,^53^ whereas its increase in GSD Ib is consistent with the accumulation of simpler structures. Notably, increased antennarity and ALN are well-recognized features of tumor glycans.^54,55^ However, in our cohort, these alterations were consistently present in all GSD Ia patients, regardless of HCA/HCC status. This suggests that they likely represent a primary metabolic effect of GSD Ia rather than a tumor-specific signature. Whether such changes might still contribute to a pro-tumorigenic environment, and thereby act as a potential risk factor, remains an open question that warrants further study.

Fucosylation also diverged between subtypes, with GSD Ia showing reduced TF, and GSD Ib exhibiting increased levels. Clinically, elevated levels of fucosylated *N*-glycans on individual proteins are established markers of HCC, exemplified by AFP-L3, a fucosylated variant of alpha-fetoprotein (AFP), ^56, 57^ and the trifucosylated tetra-antennary *N*-glycan (H7N6F3) of alpha-1 acid glycoprotein (AGP) in serum, both linked to tumor progression. ^58^ However, in our study, the fucosylation patterns were analyzed on *N*-glycans released from all proteins in patient serum/plasma, rather than from a specific one. Whether circulating total fucosylation increases or decreases in tumor patients remains debated. Increased levels of total fucosylated *N*-glycans have been observed in the serum of patients with HCC in several studies,^59–61^ whereas reduced levels of fucosylated *N*-glycans have also been reported, for example, in the serum of patients with glioblastoma and meningioma relative to healthy individuals.^62^ It will be interesting to include GSD Ia in detailed follow-up studies on the role of fucosylation in tumor development, due to the unique metabolic trigger for the early development of the tumors in this disease.

To sum up the previous findings, the identified potential glycosylation risk markers for tumor development described above, highlights hypersialylation and ALN-related alterations in GSD Ia, both of which have been previously associated with tumor developments and may hold prognostic potential. In contrast, the reduction of galactosylated diantennary *N*-glycans showed a general effect across both GSD I subtypes, and fucosylation showed a decrease overall GSD Ia, rather than the increase typically reported in classical tumor settings. These findings suggest that while some glycomic alterations in GSD Ia overlap with known tumor-associated features, while others reflect general disease effects or potentially unique pathways of tumor development in this metabolic disease.

To investigate tumor-associated glycosylation in HCA/HCC, in GSD Ia patients who developed HCA/HCC, we observed a distinct subset of differentially regulated *N*-glycans compared with the potential changes related to tumor risk hypothesized earlier. We identified four derived traits in the unique subset of GSD Ia patients with HCA/HCC, that distinguished them from both controls and non-tumor GSD Ia. Tumor-associated profiles were characterized by a shift toward less complex, oligomannose structures and increased α2,3-sialylation of fucosylated diantennary *N*-glycans. Specific studies on HCA/HCC in GSD Ia are lacking, but these alterations are consistent with tumor-associated glycosylation reported in other cancers: oligomannose *N*-glycans have been observed in breast cancer serum and on tumor cell surfaces,^63,64^ while increased α2,3-sialylation and fucosylation of diantennary *N*-glycans have been linked to pancreatic cancer progression.^65^ These parallels support the notion that the glycomic changes we observed in GSD Ia with HCA/HCC are not incidental but reflect broader mechanisms of tumor biology. Notably, the opposite regulation of α2,3-sialylation and oligomannosidic structures in GSD Ia patients with and without tumors may provide novel insight into how glycosylation dynamics mark the transition from metabolic disease to malignancy. These observations could also help with a potential distinction between biomarkers for prognosis as well as monitoring of the HCA/HCC complications in GSD Ia, though longitudinal studies would be required for the validation of our generated hypotheses.

Furthermore, we identified individual *N*-glycans as potential biomarkers. The strong discriminatory performance of individual *N-*glycans highlights their translational potential for GSD subtype classification and tumor surveillance. An AUC of > 0.9 on ROC analysis is generally considered excellent for biomarker performance.^66^ In GSD Ia, six ALN *N*-glycans reached this level, clearly discriminating from both controls and from GSD Ib, consistent with the observed overall increase of ALN. Increased ALN is a well-recognized hallmark of tumor glycosylation, ^54,55^ further supporting the biological relevance of this finding. In GSD Ib, two simple *N*-glycan compositions emerged as potential biomarkers, in line with the overall simplification of *N*-glycan structures in this subtype. In the tumor subgroup (GSD Ia), one increased oligomannosidic *N*-glycan and two decreased bisected *N*-glycans were consistent with the derived trait changes (increased TM and decreased TC), a pattern also reported in colorectal cancer.^67^ Taken together, these results indicate that individual *N*-glycans (or combinations thereof) hold considerable promise as biomarkers for GSD I stratification and tumor risk, in line with established clinical examples such as AFP-L3 for HCC.^56,57^ Future studies in independent and longitudinal cohorts will be essential to validate these candidates and assess their additive value alongside established clinical markers.

While this study represents the largest glycomic analysis of GSD I to date, some limitations should be acknowledged. Due to the rarity of the disease, the sample size was modest, particularly for the GSD Ib (n=8) and tumor (n=8, with only one HCC) subgroups. Although ROC-based performance was excellent, these small numbers raise the risk of overfitting and underscore the need for validation in larger, independent cohorts. In addition, our analyses were based on normalized signal intensities rather than absolute quantification, which may overestimate biomarker performance. Therefore, targeted quantification using glycan standards will be essential for validation. The retrospective design also meant working with both serum and plasma samples. Given the rarity and precious nature of these samples, and the absence of bias in PCA, analyzing them together was considered justified. Finally, mechanistic interpretation was constrained by incomplete proteomic coverage of glycosylation enzymes, particularly higher-order branching enzymes such as MGAT4/5, in our earlier liver tissue proteomics study.^14^

Looking forward, mechanistic insights could be strengthened through targeted proteomics or Western blotting to validate protein abundance and enzymatic activities of key glycosylation enzymes, complemented by glycoproteomics to assess site-specific glycan changes on individual proteins. In parallel, larger prospective studies will be critical to validated the biomarker potential of specific *N-*glycans, and determine their prognostic utility, particularly in monitoring the development of HCA/HCC in GSD Ia.

This study represents the first comprehensive analysis of *N*-glycosylation alterations in GSD Ia and Ib, uniquely including patients with HCA/HCC. It links impaired carbohydrate metabolism to systemic *N*-glycan remodeling, revealing the common and subtype-specific alterations in both GSD I subtypes, as well as tumor-specific changes in GSD Ia patients with tumors. In addition, the results of this study highlight *N*-glycans as promising biomarker candidates for patient stratification and the prognosis of adenomas/carcinomas in GSD I. Finally, this pioneering work provides a valuable foundation for extending glycomic investigations to other GSD subtypes and for the translational application of glycomic profiling in rare metabolic diseases.

## 4. Methods

### 4.1 Samples

This study was conducted in accordance with the principles of the Declaration of Helsinki.^68^ Blood (41 serum and 5 plasma) samples were from 17 GSD Ia patients, 8 GSD Ib patients, and 21 sex-, age-, and ethnicity-matched, healthy individuals without metabolic disorders (see **Supplementary Information, Table S7** for detailed patient characteristics). The age range of participants was 12-49 years for the GSD Ia group, 11-19 years for the GSD Ib group, and 10-50 years for the healthy controls. Notably, for patients diagnosed with HCA or HCC, blood samples were collected after confirmation of the tumor diagnosis and prior to surgical intervention. Clinical records were carefully reviewed to verify the exact timing of blood collection relative to the date of diagnosis and, when applicable, the date of surgery.

### 4.2 Sample preparation

#### 4.2.1 In solution N-glycan release

*N*-glycans were released from 2 μL plasma/serum samples (GSD Ia: n=17, GSD Ib: n=8, control: n=21). As positive control, 2 µL of a standard plasma sample (TPNG, n=5, IQ products, pooled K3EDTA plasma from healthy volunteers) was used and 2 µL H_2_O was used as negative control for the in-solution *N*-glycan release. To verify the *N*-Glycan composition with fragmentation, 2 µL original sample was pooled together for each experimental group (GSD Ia: n=1, GSD Ib: n=1, and control: n=1). Samples were mixed with 4 μL of 2% sodium dodecyl sulfate (SDS, Sigma Aldrich, Germany, cat. nr. L4390), following by 5 min shaking at 450 rpm and incubated at 60℃ for 10 min. Subsequently, 4 μL release mixture was added, which consisted of 2 μL of 4 % Nonidet P-40 substitute (NP40, Merck KGaA, Germany, cat. nr. 492016), 2 μL acidic 5 x phosphate-buffered saline solution (PBS, PH=6.0-6.5), and 0.2 μL PNGase F (1 Units/μL Roche Diagnostics GmbH, Germany, cat. nr. 11365177001). The acidic 5x PBS was prepared in house and contained 160 mM disodium hydrogen phosphate dihydrate (Sigma Aldrich, Germany, 71643), 720 mM sodium chloride (Sigma Aldrich, Steinheim, cat. nr. 746398), 20 mM potassium dihydrogen phosphate (Merck KGaA, Germany, cat. nr. 1.04872) and 85% phosphoric acid (Sigma Aldrich, Germany, cat. nr. 345245). The samples were incubated at 37℃ for 17 h.

#### 4.2.2 Sialic acid neutralization

To stabilize and neutralize the sialic acids, a sialic acid linkage-specific derivatization procedure was performed using ethyl esterification according to a previously described method.^27,69^ Briefly, 1 μL of the released *N*-glycans was transferred to 96 well PCR plates. To this, 20 μL ethyl esterification reagent comprising 250 mM 1-ethyl-3-(3-dimethylaminopropyl) carbodiimide (EDC, Thermo Scientific, U.S.A., cat. nr. 22980) and 250 mM 1-hydroxybenzotriazole hydrate (HOBt, Sigma-Aldrich, Germany, cat. nr. 54802) in ethanol absolute (EtOH, BioSolve BV, Netherlands, cat. nr. 052541) was added. The reaction mixture was incubated at 37℃ for 30 min. Subsequently, 4 μL of 25% ammonia (NH_4_OH, Merck KGaA, Germany, cat. nr. 1336-21-6) was added for amidation and then incubated again at 37℃ for 30 min. After incubation, 24 μL acetonitrile (MeCN, BioSolve BV, The Netherlands, cat. nr. 012078) was added to the mixture.

#### 4.2.3 Purification of N-glycans

*N*-glycan purification was performed by cotton HILIC SPE as described before.^27,69,70^ Briefly, approximately 2 strands of 3 mm long cotton rope (8 stranded) (Pipoos, The Netherlands) was assembled in a 20-200 µL Sapphire tip (Greiner Bio-One, The Netherlands, cat. nr. 775350) by applying air pressure (50 kPa) on top of the tip. The cotton was equilibrated using 20 μL LC-MS ultrapure water (H_2_O, BioSolve BV, The Netherlands, cat. nr. 232141) followed by 20 μL 85% MeCN for three times. After equilibration, samples were loaded on the cotton by gently pipetting the sample up and down for 20 times. Subsequently, the cotton was washed three times with 20 μL 85% MeCN containing 1% trifluoroacetic acid (*v/v*; TFA, Sigma-Aldrich, Germany, cat. nr. 302031) and 20 μL 85% MeCN. After the washing steps, samples were eluted with 7 μL ultrapure water in a new 96-well PCR plate and dried in an Eppendorf Concentrator at 60℃ for 10 min.

#### 4.2.4 Permanent cationic labelling of N-glycan reducing end

Released and purified *N*-glycans were labeled by Girard’s Reagent P (GirP, TCI Development Co. Ltd, Japan, cat. nr. G0030) based on a previous publication.^27^ The HILIC-purified *N*-glycan samples were redissolved in 2 μL of 50 mM GirP, prepared in 90% EtOH, 10% glacial acetic acid (HAc; *v/v*; Sigma-Aldrich, Germany, cat. nr. 695092). The plate was mixed at 450 rpm for 5 min and incubated at 60℃ for 1h. Following incubation, the samples were dried in the Eppendorf Concentrator at 60℃ for 10 min and subsequently dissolved in 7 μL leading electrolyte buffer (final concentration of 400 mM ammonium acetate with a pH of 2.9; Sigma-Aldrich, Germany, cat. nr. A2706) for CE-ESI-MS analysis.

### 4.3 Capillary electrophoresis (CE)

The CE separation was performed as described previously.^27^ In brief, the experiments were conducted on a CESI 8000 plus system (Sciex, Framingham, U.S.A.) with a 91 cm long bare-fused silica capillary (30 μm internal diameter and 150 μm outer diameter; cat. nr. SN20752605). Before use, the capillary was conditioned by immersing the spray tip in MeOH (BioSolve BV, Netherlands, cat. nr. 136841), while the separation and conductive lines were rinsed with MeOH at 100 psi for 10 min and 3 min, respectively. Subsequently, the tip was immersed in H_2_O, and the separation line was rinsed at 100 psi for 10 min consecutively with H_2_O, 0.1 M NaOH (Sigma-Aldrich, Steinheim, Germany, cat. nr. 720698-100ML), 0.1 M HCl (Sigma-Aldrich, Steinheim, Germany, cat. nr. H9892-100ML), H_2_O and the background electrolyte (BGE; 20% HAc). This was followed by a final rinse of the conductive line with BGE at 100 psi for 3 min. Before each analysis, the capillary was rinsed consecutively at 100 psi with 0.1 M NaOH (2.5 min), 0.1M HCl (3 min), H_2_O (4 min), and BGE (4 min) while the conductive line was rinsed with BGE for 3 min at 75 psi. The samples were stored in the system at 10 °C. Sample injection was performed hydrodynamically by applying 10 psi for 60 s, which corresponded to 13.6% of the total capillary volume (86 nL). Following each injection, a post plug of BGE was injected by applying 0.5 psi for 25 s, accounting for 0.3% of the capillary volume. The analysis was conducted by maintaining a constant flow with 0.5 psi pressure and 20 kV over the capillary with the temperature set to 20°C.

### 4.4 Mass spectrometry (MS)

The MS method followed a previously described procedure.^27^ Briefly, the CESI 8000 system was connected to a Maxis Plus quadrupole time-of-flight-(qTOF-) MS (Bruker Daltonics, Germany) via a sheathless CE-ESI-MS interface (Sciex, U.S.A.), ensuring optimal alignment between the capillary spray tip and the front of the nanospray shield (Bruker Daltonics, Germany). All experiments were performed in positive-ionization mode, with a stable electrospray maintained by establishing an electric field between the CE (ground potential) and a negatively charged spray shield (between - 1100 and -1300 V). The drying gas temperature and flow rate were set at 150°C and 1.2 L/min, respectively. To minimize the in-source decay, the collision cell energy and the quadrupole ion energy were set at 5.0 eV, and the pre-pulse storage was set at 25.0 µs. To enhance ionization, a dopant enriched nitrogen (DEN) gas was implemented,^71^ MeCN was used as dopant at 0.2 bar. Fragmentation experiments were performed only on the pooled samples at a frequency of 1.00 Hz, targeting the three most abundant precursor ions within *m/z* 100–2200 range. An absolute threshold of 1822 counts was applied, with active exclusion settings to exclude after two spectra and release after one min, and reconsider precursor if the current intensity exceeded five times the previous intensity. Precursor ions were isolated with a width of 8-15 Thomson, depending on their *m/z* values. Collision energies were applied in a linear *m/z* dependent manner, ranging from 20 eV at m/z 500 to 70 eV at m/z 2000 for all charge states (1-3). To facilitate the identification of low-abundant *N*-glycans, an exclusion list was applied (**Supplementary Information, Table S8**) to prevent repeated selection of precursor ions already subjected to previous MS/MS experiments (width ± 0.5).

### 4.5 Data analysis and statistics

Raw mass spectrometric data were processed using GlycoGenius v. 1.0.7. ^72^ An accuracy level of 30 ppm was applied throughout the analysis. Data curation was carried out with the following parameters: a minimum isotopic fitting score of 0.8, curve fitting score of 0.9, and a signal-to-noise (S/N) ratio threshold of 3. The mass accuracy tolerance was set to ±10 ppm, and the reducing end tag was specified as GirP, with amidation or ethyl esterification enabled. The retention/migration time range for analysis was set from 30 to 60 minutes. Data processing was performed on a server utilizing 24 central processing unit (CPU) cores. For glycan library generation, the following compositional constraints were applied: total monosaccharides ranging from 5 to 22, deoxyhexose (dHex) from 0 to 2, sialic acids (Neu5Ac) from 0 to 4, hexoses (Hex) from 3 to 10, and *N*-acetylhexosamines (HexNAc) from 2 to 8. The information of all raw data and associated supplements from this study can be found in the section **Data sharing statement**.

*N*-glycans identified in more than 50% of samples within at least one study group were retained for analysis. Missing values were imputed using MetaboAnalyst (version 6.0) by replacing one-fifth of the minimum positive value for the corresponding variable. Subsequent data analysis was performed in RStudio (version 4.2.2). For each *N*-glycan, the percentage abundance, group mean, and relative standard deviation (RSD) were calculated. Log2^Fold^ ^Change^ (Log2^FC^) was computed for pairwise group comparison, alongside derived glycan traits. Statistical significance was assessed using p-values and false discovery rates (FDR), calculated in Microsoft Excel. *N*-glycans with FDR ≤ 0.05 were considered significantly altered. Data visualizations were carried out using R packages, including principal component analysis (PCA) and volcano plots by “ggplot2”, box plots and violin plots by “ggpubr”, heatmap by “pheatmap”, receiver operating characteristic (ROC) curve by “pROC”.

### 4.6 Glycosylation-related enzymes intensity measurement

To investigate the mechanisms of altered *N*-glycan patterns in GSD I, we explored the intensity of 42 key glycosylation-related enzymes, which were related to sialylation, branching, poly-*N*-acetyllactosamine (ALN), and fucosylation (**Supplementary Information, Tables S6**) in our previous untargeted proteomics screenings of these patient samples.^14^ Briefly, serum/plasma samples from the same cohort and 14 liver samples (2 GSD Ia without HCA/HCC, 1 GSD Ia HCC tissue, 1 GSD Ib, and 10 controls) were analyzed by untargeted proteomics. Notably, a pair of liver tissues, HCC and non-HCC, was collected from tumor and non-tumor adjacent tissues from the same patient. We selected glycosylation-related protein values from both serum/plasma and liver datasets. As only one GSD Ib liver sample was available, proteins were included if they exhibited > 50% change in GSD Ib relative to the mean of controls and GSD Ia, while those with values overlapping at least one control were excluded. HCC samples (L_A02_HCC) were not analyzed separately, as their protein profiles were highly correlated with the adjacent non-tumor liver tissue from the same patient (L_A02). Proteomics data were analyzed retrospectively and, although not included in the primary results, were discussed to provide additional context for the interpretation.

## Acknowledgements

The authors gratefully acknowledge the support from the Chinese Scholarship Council (CSC) for Ruiqi Xiao (202106220094), Coordenação de Aperfeiçoamento de Pessoal de Nível Superior (CAPES), awarded to Hector F. B. R. Loponte and the Dutch Research Council (NWO) that funded the X-omics Road Map program (project 184.034.019) and project VI.Veni.222.262, awarded to Guinevere S. M. Lageveen-Kammeijer The authors are grateful for the opportunities and support afforded by the Sector Plan Pharmaceutical Sciences, which was implemented in the overarching Sector Plan Beta II, put into action by the Dutch Ministry of Education, Culture and Science (OCW). The authors are grateful to Maaike H. Oosterveer for collecting serum and plasma samples from patients and controls.

## 6. List of abbreviations

A: antennarity
ALN: LacNAc; poly-*N*-acetyllactosamine
Am: α2,3-sialylation
B: bisection
BGE: background electrolyte
C: complex
CAPES: Coordenação de Aperfeiçoamento de Pessoal de Nível Superior
CDG: congenital disorders of glycosylation
CE-ESI-MS: capillary electrophoresis-electrospray ionization mass spectrometry
CPU: central processing unit
CSC: Chinese Scholarship Council
DEN: dopant enriched nitrogen
dHex: deoxyhexose
E: α2,6-sialylation
EDC: 1-ethyl-3-(3-dimethylaminopropyl) carbodiimide
ER: endoplasmic reticulum
EtOH: ethanol absolute
Fru: fucose
F: fucosylation
Fa: antennary fucosylation
Fc: core fucosylation
FDR: false discovery rates
Fru: fructose
G: galactosylation
G6P: glucose 6-phosphate
G6PC1/G6PC: glucose-6-phosphatase
G6PT: glucose-6-phosphate transporter
Gal: galactose
GalNAc: *N*-acetylgalactosamine
GirP: Girard’s Reagent P
Glc: glucose
GlcN: glucosamine
GlcNAc: *N*-acetylglucosamine
GSD: glycogen storage disease
H: Hex hexose (Gal; Man; Glu)
Hy: hybrid
HAc: acetic acid
HCA: hepatocellular adenoma
HCC: hepatocellular carcinoma
HCl: hydrochloric acid
HOBt: hydroxybenzotriazole hydrate
Kdn: keto-deoxy-nonuronic acid
M: oligo-mannosidic
Man: mannose
MAN1A1: mannosidase α class 1A member 1
MAN1A2: mannosidase α class 1A member 2
ManNAc: *N*-acetylmannosamine
MeCN: methyl cyanide
MeOH: methanol
MGAT1: mannosyl (α-1,3-)-glycoprotein β-1,2-N-acetylglucosaminyltransferase 1
MGAT2: mannosyl (α-1,6-)-glycoprotein β-1,2-N-acetylglucosaminyltransferase 2
MS: mass spectrometry
N; HexNAc: *N*-acetylhexosamine (GalNAc; ManNAc; GlcNAc)
NaOH: sodium hydroxide
Neu5Ac: sialic acids
Neu5Ac; S: *N*-acetyl-neuraminic acid
Neu5Gc: *N*-glycolyl-neuraminic acid
NH4OH: ammonia
NP-40: nonidet P-40 substitute
NWO: Dutch Research Council
OCW: Dutch Ministry of Education, Culture and Science
P: pauci-mannosidic
PBS: phosphate-buffered saline
PCA: principal component analysis
qTOF-MS: quadrupole time-of-flight mass spectrometry
ROC: receiver operating characteristic
RSD: relative standard deviation
S: sialylation
ST6GAL1: ST6 β-galactoside α-2,6-sialyltransferase 1
S/N: serial number
TC: total complex
TF: total fucosylation
TFA: trifluoroacetic acid
TG: triglycerides
TM: total oligomannosidic
TP: total pauci-mannosidic
TS: total sialylation
TU: total unclassified
U: unclassified
UDP-Gal: uridine diphosphate-galactose
UMCG: University Medical Center Groningen

