## Supplementary figures for "Decoding the Glycan Signature: Unraveling *N*-Glycosylation Alterations in Glycogen Storage Disease Ia and Ib"

* Shared last author

**Corresponding Authors:** Justina C. Wolters, Guinevere S. M. Lageveen-Kammeijer | Email address: | Address, Antonius Deusinglaan 1, 9713 AV Groningen, The Netherlands | Internal Zip Code: XB20,, P.O. Box 196, 9700 AD Groningen

### TABLE OF CONTENTS

**S-1. Supplementary Figures – 1 – 8 ......... 2**

### S-1. SUPPLEMENTARY FIGURES – 1 – 8

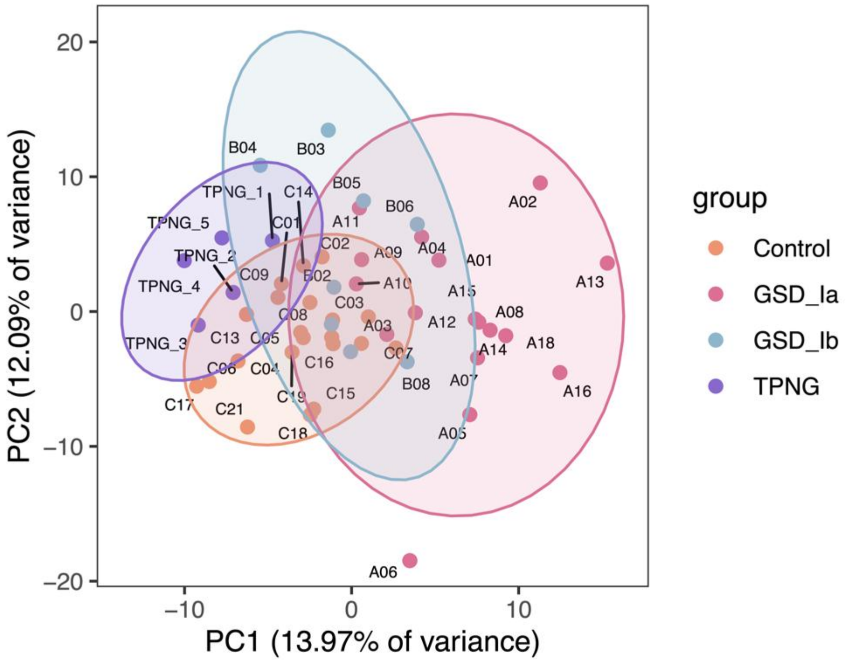


**Figure S1. The Principal Component Analysis (PCA) plot illustrates the proteomic profile separation among groups and standard plasma replicates.** In the cohort, a total of 21 controls, 17 GSD Ia, 8 GSD Ib samples, and 5 standard plasma samples (TPNG) were analyzed. The cluster of five TPNG indicated minimal technical variation in this experiment, and the main source of variation comes from biological differences. The samples from the same patient but at different time points were labeled using 'a', 'b', and 'c' (e.g., A03a and A03b). The replicate samples from standard plasma samples were labeled using '1' to '5' (e.g., TPNG1 and TPNG2).

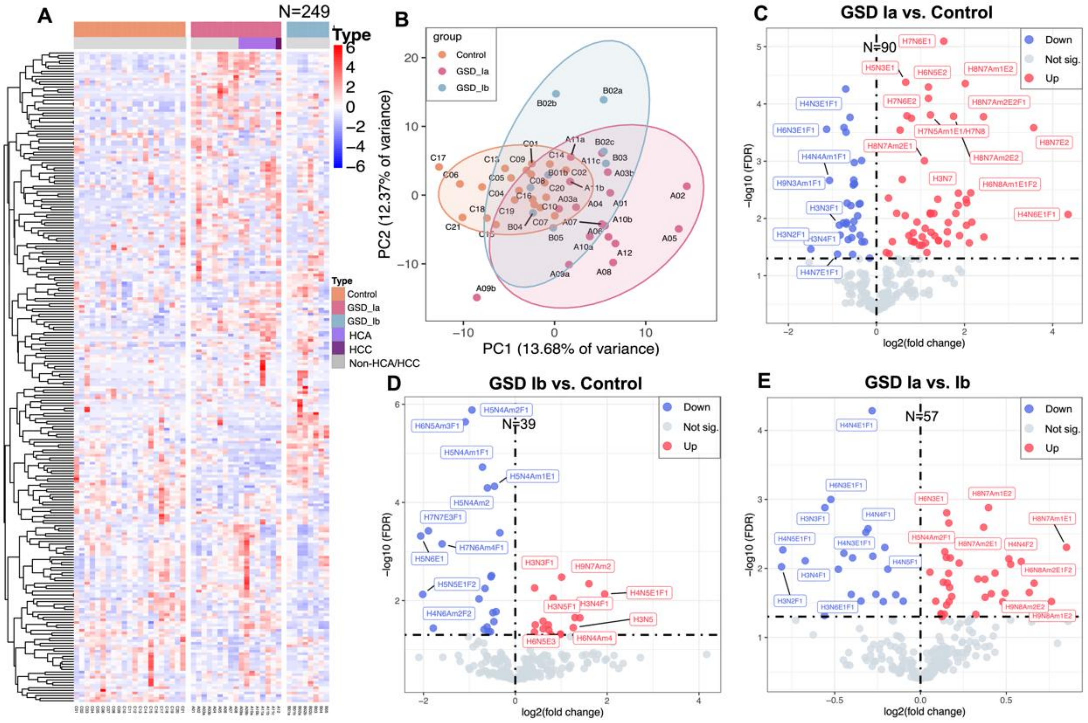


**Figure S2. Overview of the total plasma/serum *N*-glycome for each individual across control, GSD Ia, and GSD Ib groups. (A)** Heatmap showing the z-score of all *N*-glycans detected in all samples from all groups. **(B)** Principal Component Analysis (PCA) plots illustrate the separation of glycomics profiles among groups. **(C-E)** Volcano plots of differential *N*-glycans between groups. Significantly up-regulated and down-regulated *N*-glycans (FDR < 0.05) are highlighted in red and blue, respectively. "N" refers to the number of significantly changed *N*-glycans, including "up" and "down" *N*-glycans. "Not sig." refers to *N*-glycans that show no significant changes. The cohort consisted out of 21 controls, 17 GSD Ia (including 7 HCA, 1 HCC, and 9 without HCA/HCC), and 8 GSD Ib samples were grouped. The samples from the same patient but at different time points were labeled using 'a', 'b', and 'c' (e.g., A03a and A03b).

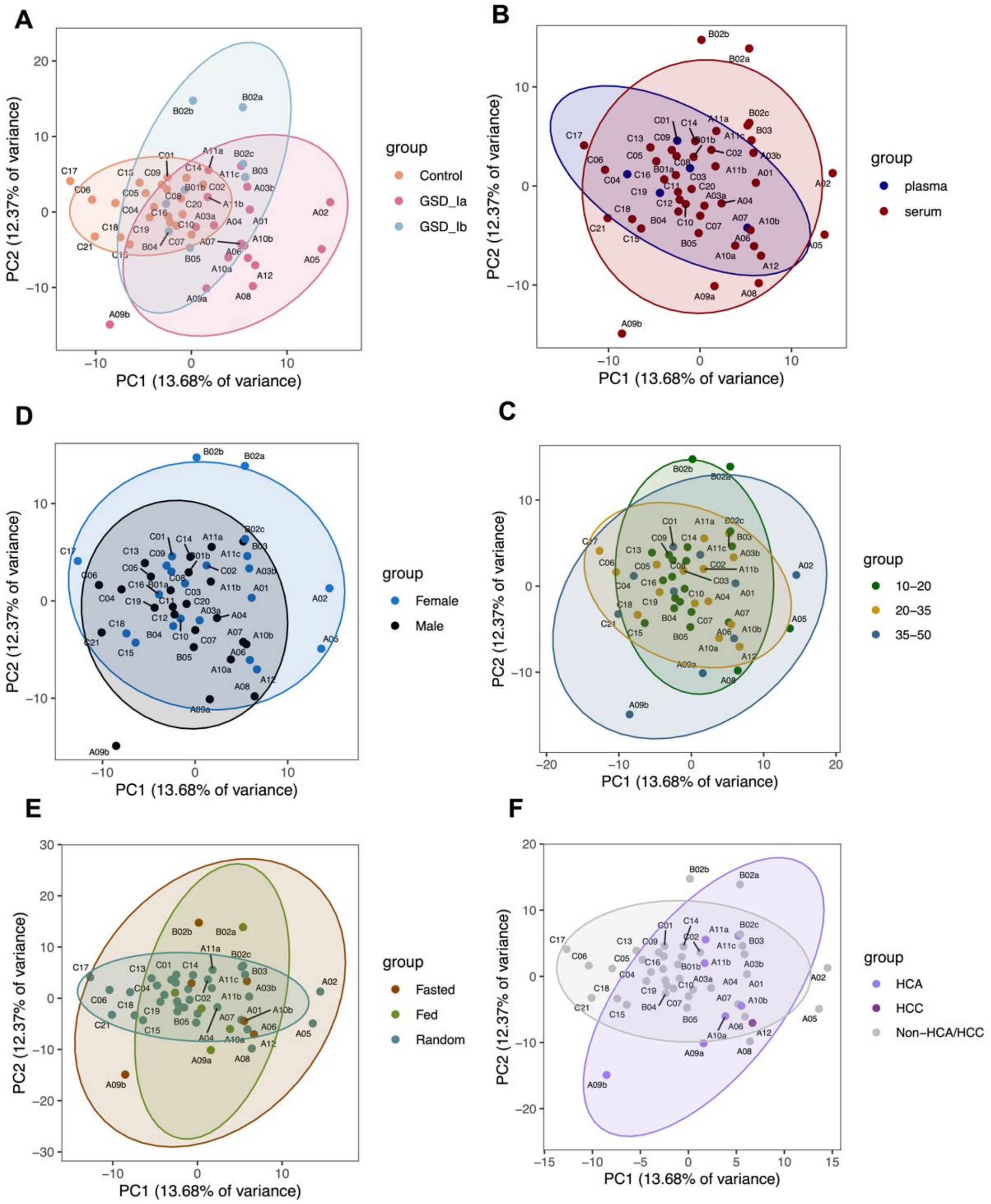


**Figure S3. The Principal Component Analysis (PCA) plot illustrates the proteomic profile separation among groups.** The cohort consisted out of 21 controls, 17 GSD Ia (including 7 HCA, 1 HCC, and 9 without HCA/HCC), and 8 GSD Ib samples. The samples from the same patient but at different time points were labeled using 'a', 'b', and 'c' (e.g., A03a and A03b).

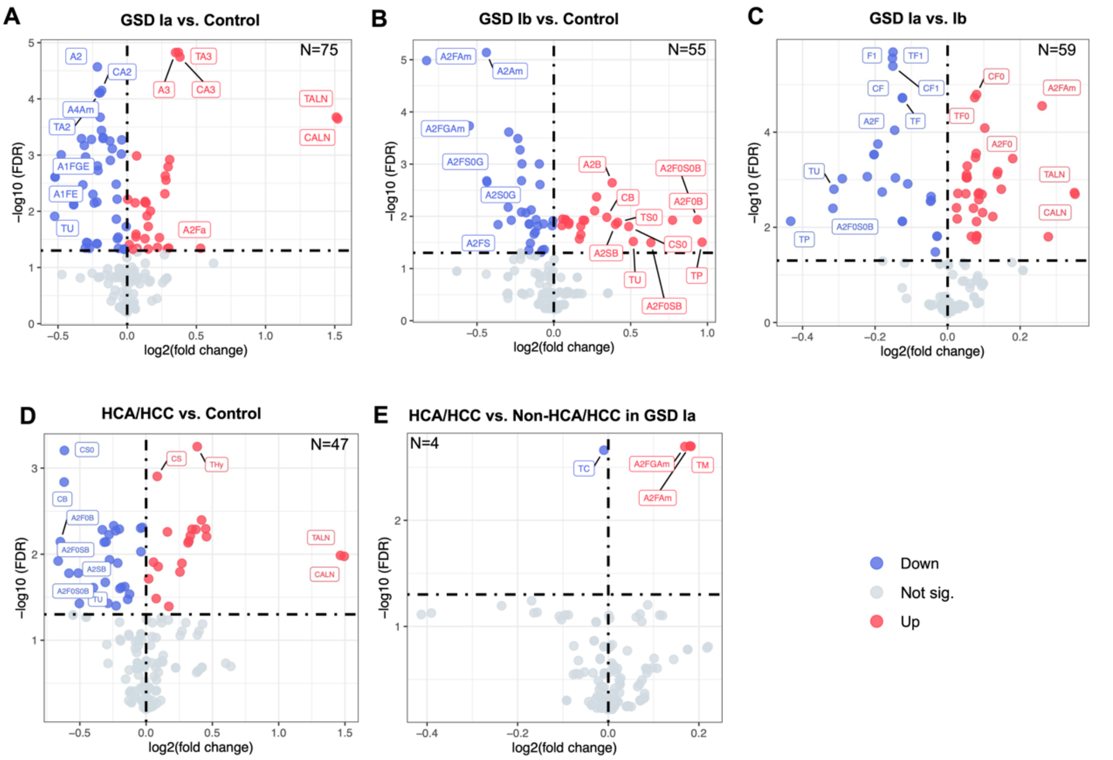


**Figure S4. Volcano plots of differential derived *N*-glycan traits** between (**A**) GSD Ia (n=17) and control (n=21), (**B**) GSD Ib (n=8) vs. control, (**C**) GSD Ia vs. Ib, (**D**) GSD Ia with hepatocellular adenoma/carcinoma (HCA/HCC) (n=8) vs. control, and (**E**) HCA/HCC vs. Non-HCA/HCC in GSD Ia (n=9). Significantly up-regulated and down-regulated *N*-glycans (FDR < 0.05) are highlighted in red and blue, respectively. 'N' refers to the number of significantly changed *N*-glycans, including 'up' and 'down' *N*-glycans. 'Not sig.' refers to *N*-glycans that show no significant changes.

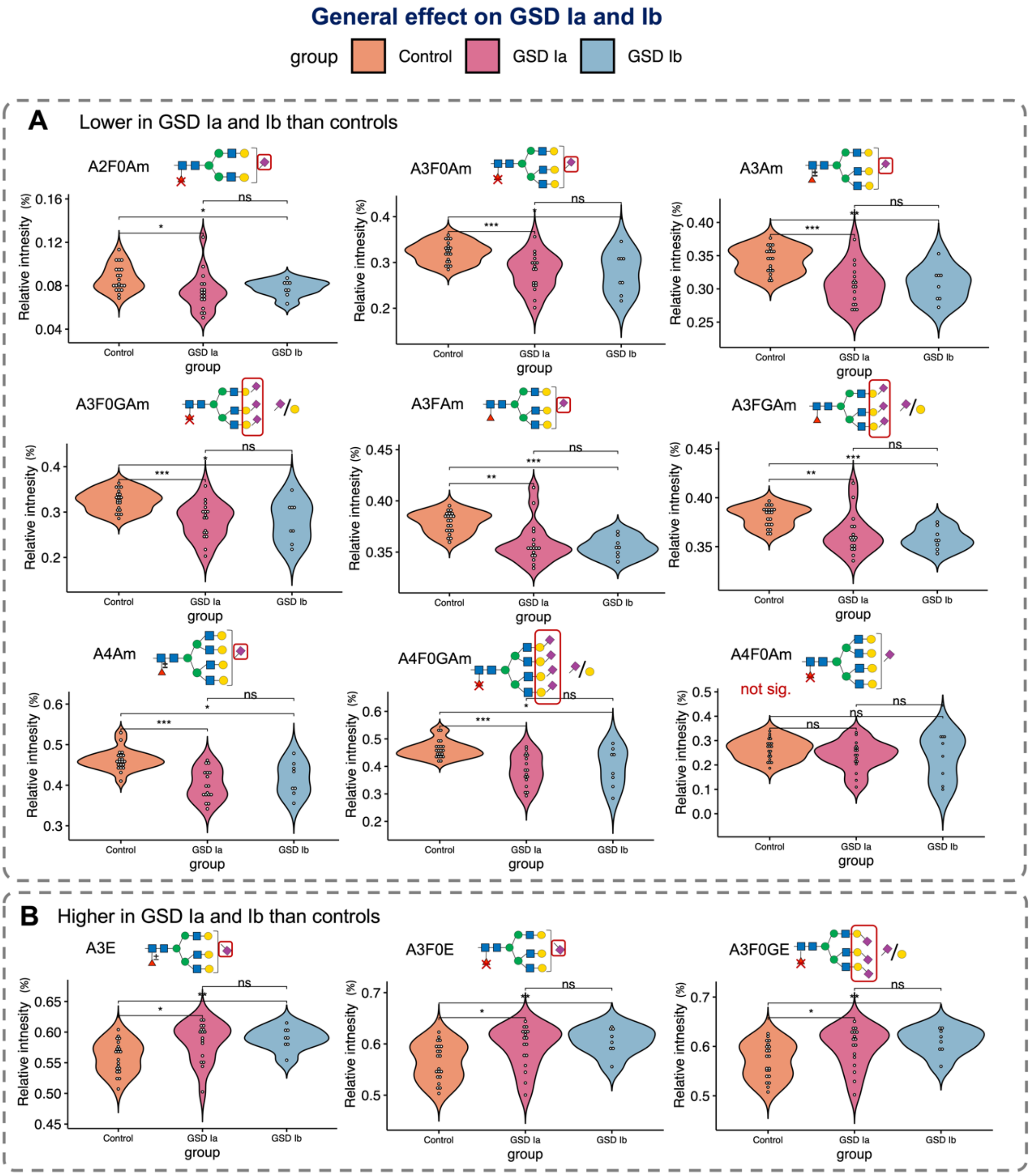


**Figure S5. Violin plots of the derived traits with the general effect on GSD Ia and Ib. (A)** Lower in GSD Ia (n=17) and Ib (n=8) than control (n=21). **(B)** Higher in GSD Ia and Ib than control. Significance levels: FDR<0.05 (*), FDR<0.01 (**), FDR<0.001 (***). "ns" refers to no significant changes. Example of the derived trait labels: "A2F0Am" refers to nonfucosylated α2,3-sialylated diantennary *N*-glycans. "A2F0GAm" refers to the ratio of α2,3-sialic acid and galactose on nonfucosylated α2,3-sialylated diantennary *N*-glycans. "A4Am" refers to α2,3-sialylated tetra-antennary *N*-glycans. "A3E" refers to α2,6-sialylated triantennary *N*-glycans.

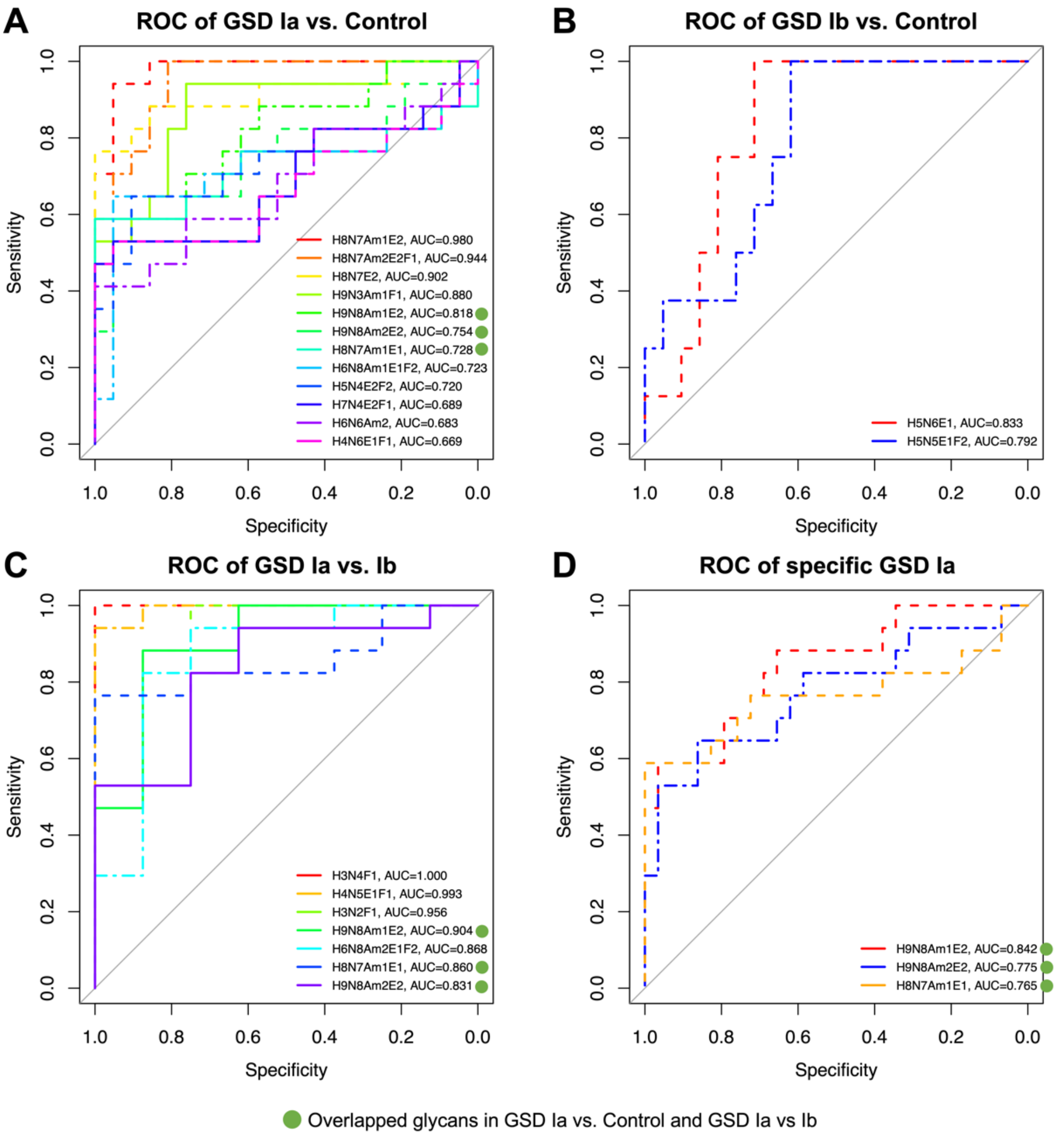


**Figure S6. ROC curves of *N*-glycan-based comparisons.** ROC curves of *N*-glycans with $\log2^{FC} > \pm 2$ in (**A**) GSD Ia (n=17) vs. control (n=21) and (**B**) GSD Ib (n=8) vs. control, and (**C**) GSD Ia vs. GSD Ib. (**D**) ROC curve of GSD Ia-specific *N*-glycans that overlap between both GSD Ia vs. control and GSD Ia vs Ib comparisons. (**E**) ROC curve of a combined model for distinguishing GSD Ia from both GSD Ib and controls, including six *N*-glycans with AUC>0.9 in GSD Ia vs. control, along with GSD Ia-specific glycans. (**F**) ROC curve of a combined model distinguishing GSD Ib from controls.

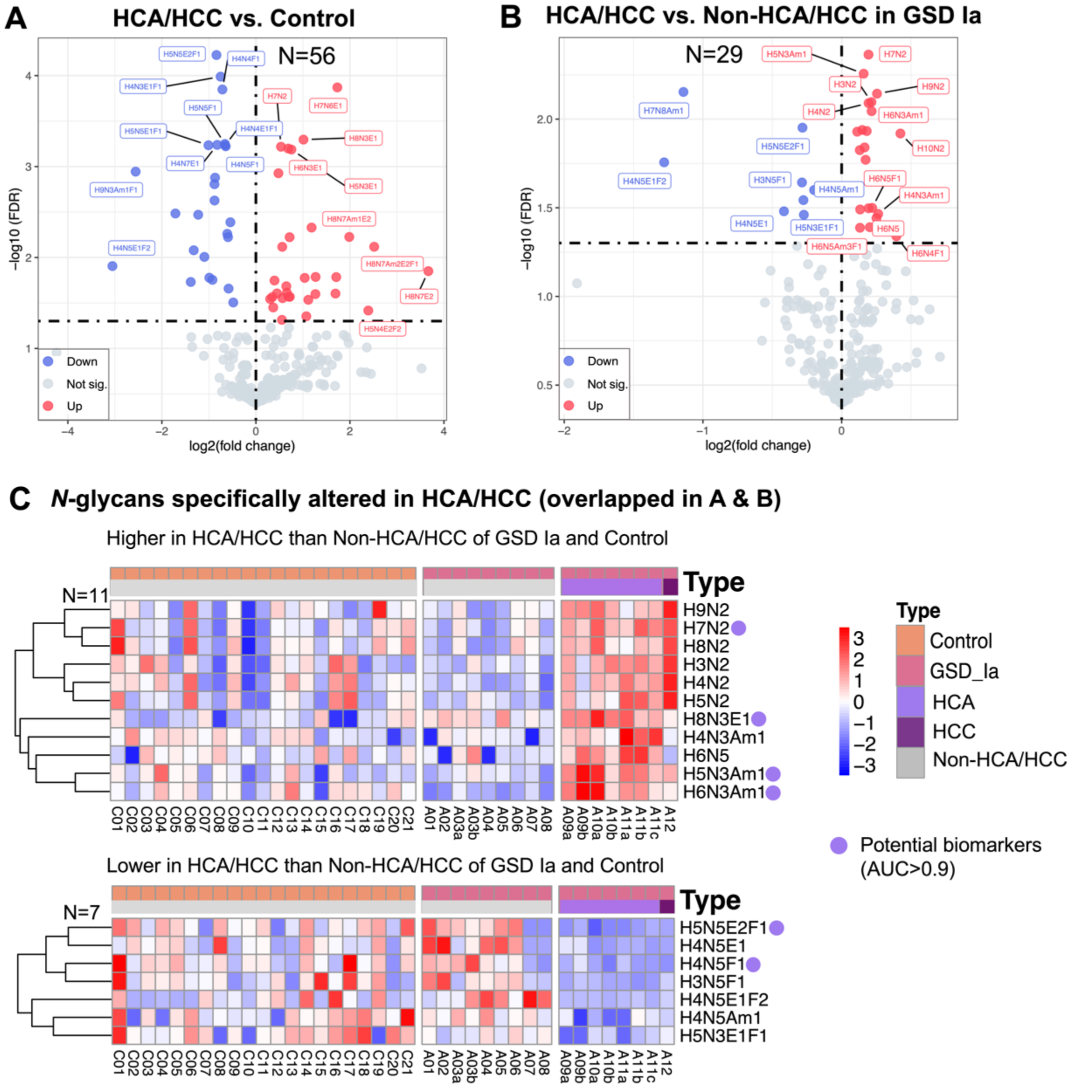


**Figure S7. Potential biomarkers for HCA/HCC in GSD Ia. Volcano plots of differential *N*-glycans** between (**A**) GSD Ia with HCA/HCC (n=8) vs. control (n=21), and (**B**) HCA/HCC vs. non-HCA/HCC in GSD Ia (n=9). (**C**) Heatmap showing the z-score of all *N*-glycans, with consistent up- or down-regulation patterns observed in A and B. Significantly up-regulated and down-regulated *N*-glycans (FDR < 0.05) are highlighted in red and blue, respectively. 'N' refers to the number of significantly changed *N*-glycans, including 'up' and 'down' *N*-glycans. 'Not sig.' refers to *N*-glycans that show no significant changes.

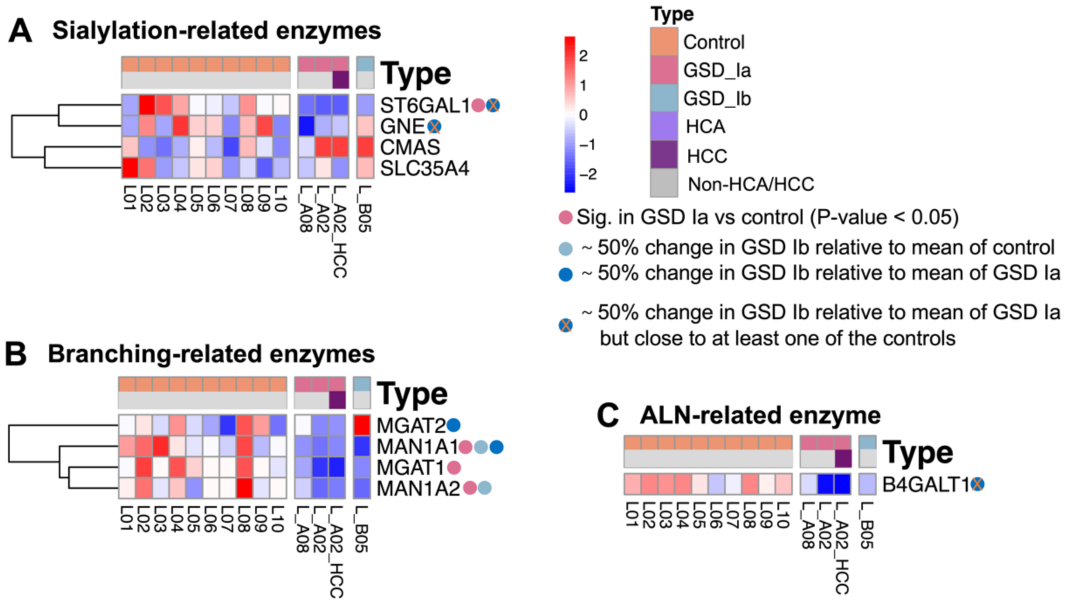


**Figure S8. Summary of potential *N*-glycan biomarkers of GSD Ia, GSD Ib, and HCA/HCC. (A)** Boxplots showing the *N*-glycans with log2$^{FC}$>2 in GSD Ia (n=17) vs. control (n=21), GSD Ib (n=8) vs. control, and GSD Ia vs. Ib. **(B)** Boxplots showing the Log2$^{FC}$ of *N*-glycans with the same changes in HCA/HCC (n=8) versus control and Non-HCA/HCC in GSD Ia (n=9). **(C)** Receiver operating characteristic (ROC) curve of *N*-glycans showing in B.
